## Supplementary Materials for "*In-silico* Optimisation of Mass Spectrometry Fragmentation Strategies in Metabolomics"

### In-silico Optimisation of Mass Spectrometry Fragmentation Strategies in Metabolomics - Supplementary Materials

#### 1 MS1 Simulations

Figure 1 shows the overall characteristics of m/z, RT and intensity values from running XCMS peak picking on the resulting simulated mzML file and 4 *multi-beer* samples for comparison (labelled *multi-beer-1*, ..., *multi-beer-4*). It can be observed from Figure 1 that all m/z and RT boxplots show similar profiles between the 4 randomly selected real beer samples and the simulated sample from ViMMS. We explain the slightly higher intensity distribution in Figure 1c due to the fact that in our sampling scheme, a chemical having high sampled maximum intensity values could be assigned a large ROI that spans a big RT range. During peak picking, these potentially-noisy ROIs are detected as multiple high-intensity peaks by XCMS, resulting in an upward shift in the intensity distribution of detected MS1 features from the simulated mzML file. Improving the regions of interest detection scheme to address this issue is a future work.

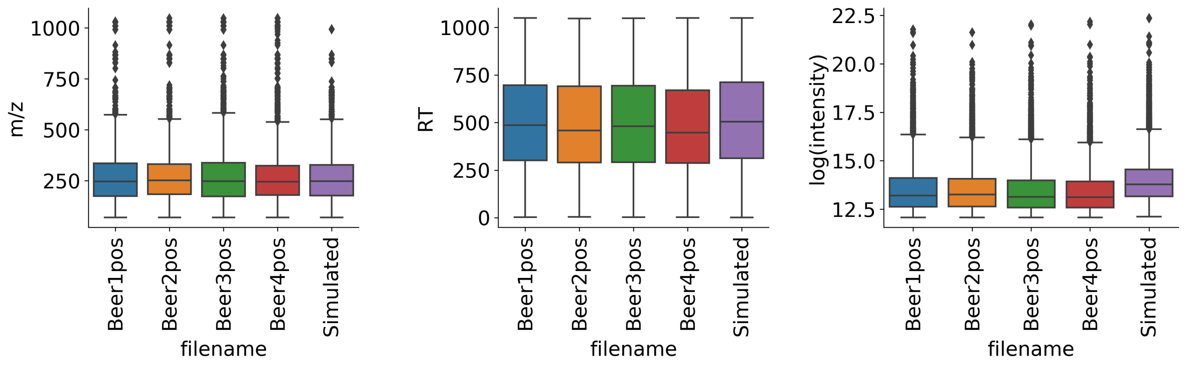


(a) m/z (b) RT (c) Log Intensity

Figure 1: A figure showing boxplots of the simulated data and some of the *multi-beer* samples.

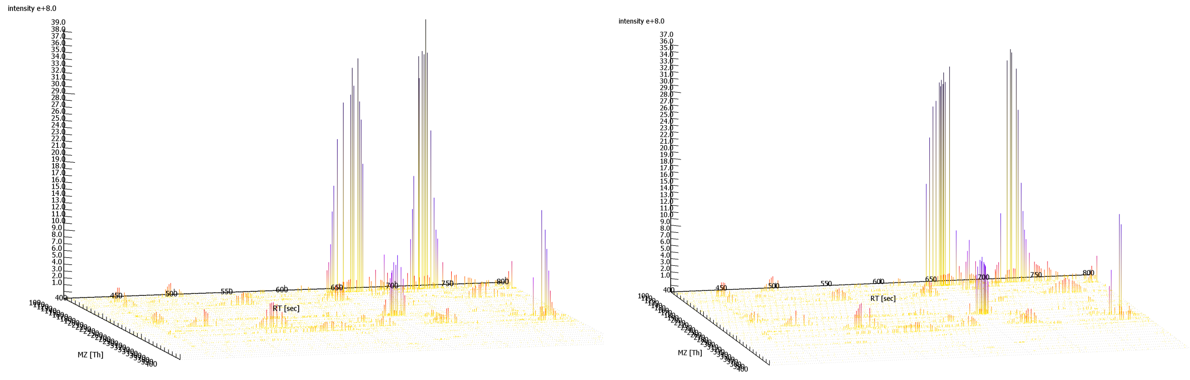


(a) Real (b) Simulated

Figure 2: ToppView visualisation of mz range (100 - 400) ppm and RT range (400 - 800)s for a Top-10 DDA fragmentation strategy in ToppView for (**a**) a *multi-beer* sample and (**b**) for the same file generated using ViMMS

### 2 Top-N Simulations

From the actual *multi-beer-1* fragmentation mzML, we extracted 512,540 ROIs across all scans using a 10 ppm tolerance that determines when peaks should be grouped under one ROI. Only ROIs with at least one peak above the minimum intensity threshold of 1.75E5 are kept. This results in 10,190 ROIs, which are converted into chemical objects in ViMMS, having their corresponding normalised chromatographic peak shapes derived from the ROIs. Simulated Top-10 DDA fragmentation was performed using the same fragmentation parameters used to generate the actual beer1 data (N=10, DEW=15s). Simulation took approximately 2 minutes in ViMMS running on an Intel Core i9 laptop. The generated mzML file and simulator state were loaded into ToppView and Jupyter Notebook for further analysis.

Visual inspection of the resulting spectra in ToppView shows that the mzML files from ViMMS and the *multi-beer-1* data look broadly similar (Figure 2). Comparing the number of scans, we obtained a total of 9,891 scans from the simulator to 9,406 scans in the actual beer mzML, resulting in 3,027 fewer MS1 features that can be picked by XCMS from the generated mzML file (Table 1).

To evaluate these differences quantitatively, we matched precursor ion information from the real *multi-beer 1* mzML to the generated mzML from ViMMS. 6,757 out of 7,655 (88%) precursor ions can be matched from the actual *multi-beer 1* file to the generated file when matching up to 2 decimal places for the m/z values and using 15s tolerance window for retention time. It can be seen from Figure 3 that fragmentation events from both the real and simulated files are in close proximity to each other.

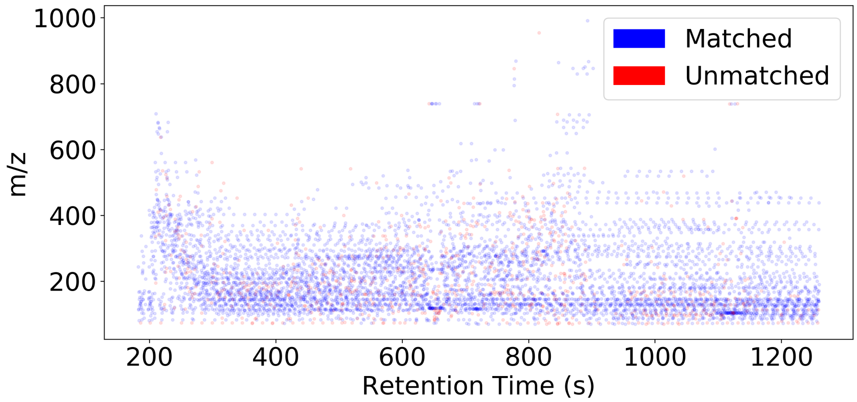


Figure 3: Locations of precursor peaks extracted from MS/MS scans in original *multi-beer-1* data that could be matched to the generated mzML data in ViMMS. The colours indicate where the MS/MS occurred in close proximity to an MS/MS in the real beer data (blue for matched fragmentation events and red for unmatched ones). Scans were performed in both cases using a Top-10 DDA fragmentation strategy.

Table 1: Table showing XCMS peak picking results on the real vs simulated files for the original beer data.

| | **Generated mzML** | **Real mzML** |
|---|---|---|
| **Number of MS1 scans** | 1,267 | 1,751 |
| **Number of MS2 scans** | 8,624 | 7,655 |
| **Number of peaks picked by XCMS** | 12,801 | 15,828 |

### 3 Alternative scenario: varying $N$s when both full-scan and Top-$N$ data are available.

Consider a scenario where within an experimental batch, full-scan and Top-N data are acquired. In a typical analysis of this scenario, the MS1 peaks detected by XCMS from the full-scan MS1 file serve as the ground truth of peaks we wish to fragment. To compute performance in this scenario, XCMS' CentWave peak detection is performed on the full-scan input files mzML files, resulting in the number of ground truth MS1 features shown in Table 2.

Table 2: Table showing the number of ground truth MS1 features from XCMS peak picking for the two full-scan (MS1-only) *multi-beer* and *multi-urine* data in the Top-N experiments.

| | **Number of ground truth MS1 features** |
|---|---|
| *multi-beer-1* | 9,198 |
| *multi-beer-2* | 9,805 |
| *multi-urine-2* | 7,188 |
| *multi-urine-3* | 7,664 |

For evaluation, we provide the following definition of positive and negative instances (illustrated in Figure 4):

**True Positives (TP)**: peaks from ground truth (found in full-scan files) that are fragmented above the minimum intensity threshold.

**False Positives (FP)**: peaks from ground truth that are not fragmented + peaks from ground truth that are fragmented below the minimum intensity threshold.

**False Negatives (FN)**: peaks not from ground truth that are fragmented above the minimum intensity threshold.

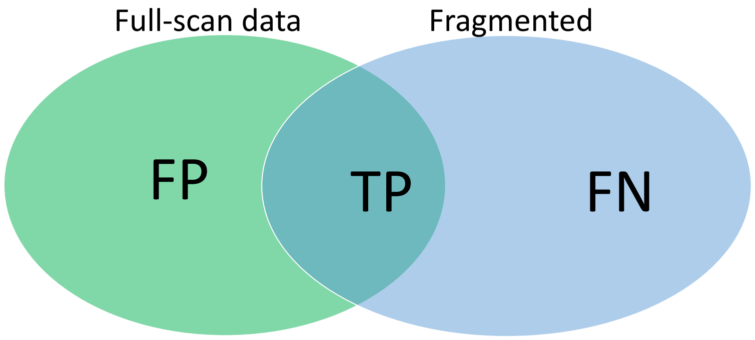


Figure 4: Definition of True Positives (TP), False Positives (FP) and False Negatives (FN) for performance evaluation of Top-N DDA fragmentation strategy (Scenario 1). The blue circle in the Venn diagram refers to all peaks that are fragmented above the minimum MS1 intensity threshold, while the green circle refers to all ground truth peaks (MS1 features) found by XCMS' CentWave from the full-scan files.

The results in this experiment shows that increasing N results in lower precision (Figure 5a), while recall increases with $N$ initially but flattens (Figure 5b). Assessing the $F_1$ score (Figure 5c), which is the harmonic average of precision and recall, we see that the best fragmentation performance (as represented by the $F_1$ score) is reached between $N$ from 10 to 20 and plateaus after 20, suggesting that no further performance gain is obtained from increasing the number of precursor ions to fragment.

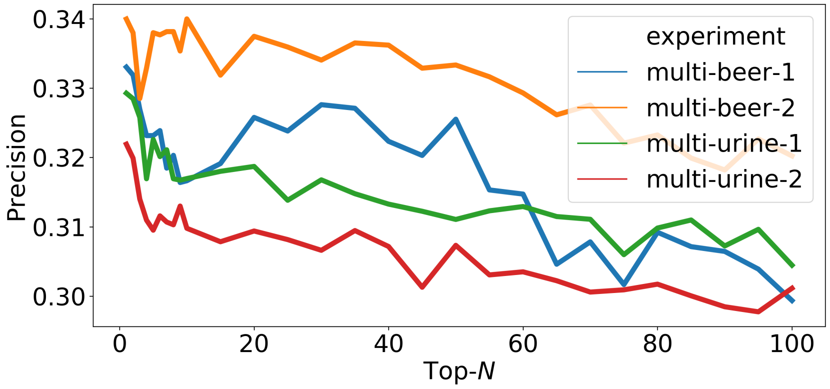


(a) Precision

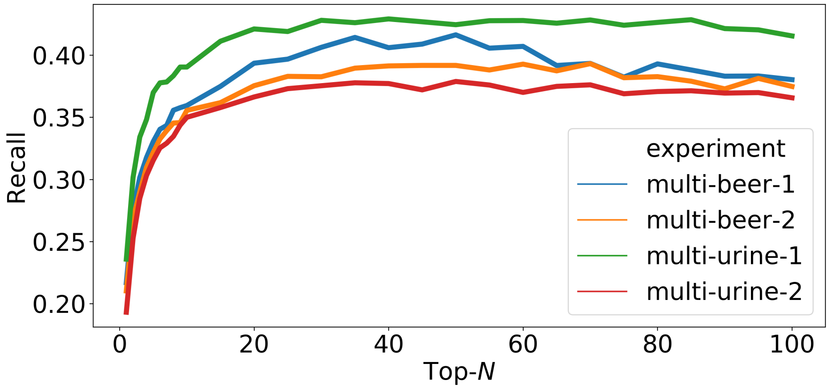


(b) Recall

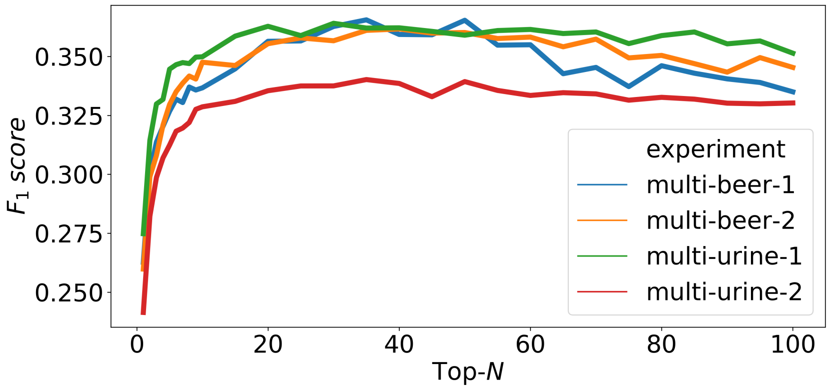


(c) $F_1$

Figure 5: Figures showing (a) precision, (b) recall and (d) $F_1$ score for peak picking performance as $N$ changes in Top-N DDA experiments in ViMMS based on the classification specifications given in Figure 4.

### 4 Varying Multiple Parameters in the Top-$N$ Simulations

Figure 6 shows boxplots comparing the $F_1$-scores (representative of fragmentation performance) of all parameter combinations from the real and simulated data. The results from our simulator generally match the results from the real data, although with a slightly greater spread in the simulated fragmentation performance. Exploring the results in detail, we see that the best fragmentation performance is obtained at $N = 20$ and $DEW = 30$ for both datasets and the worst at $N = 1$ and $DEW = (15, 30, 60)$ (Table 3). We explain our findings by the fact that the best performance is obtained at the parameter combinations with the lowest trade-off between fragmentation performance and MS2 peak picking quality. The poor results came about from when both $N$ and $DEW$ are too small. In the former, not enough unique MS1 peaks are fragmented due to dynamic exclusion effect, whereas in the latter case, the quality of peak picking from fragmentation files decrease significantly, affecting the number of true positives obtained.

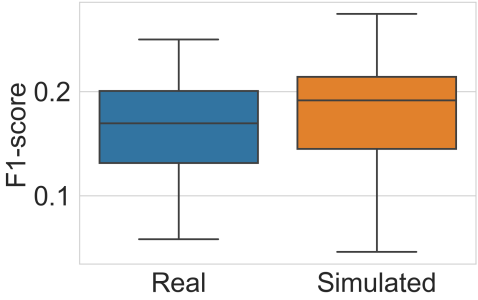


Figure 6: Distributions of F1-scores for all parameter combinations for the real and simulated *BeerQCB* data.

Table 3: The highest and lowest fragmentation performance for real and simulated results.

| Rank | Real | | | | | Simulated | | | | |
|---|---|---|---|---|---|---|---|---|---|---|
| | N | DEW | Prec | Rec | F1 | N | DEW | Prec | Rec | F1 |
| **1** | 20 | 30 | 0.820 | 0.147 | 0.250 | 20 | 30 | 0.799 | 0.166 | 0.274 |
| **2** | 10 | 30 | 0.722 | 0.135 | 0.228 | 20 | 60 | 0.761 | 0.158 | 0.262 |
| **3** | 15 | 30 | 0.769 | 0.132 | 0.226 | 15 | 30 | 0.739 | 0.157 | 0.258 |
| **-1** | 1 | 15 | 0.771 | 0.030 | 0.058 | 1 | 15 | 0.793 | 0.024 | 0.046 |
| **-2** | 1 | 30 | 0.757 | 0.039 | 0.074 | 1 | 30 | 0.743 | 0.033 | 0.062 |
| **-3** | 1 | 60 | 0.738 | 0.048 | 0.091 | 1 | 60 | 0.741 | 0.046 | 0.087 |
